# Patterns of linkage disequilibrium reveal genome architecture in chum salmon

**DOI:** 10.1101/729574

**Authors:** Garrett McKinney, Megan V. McPhee, Carita Pascal, James E. Seeb, Lisa W. Seeb

## Abstract

Many studies exclude loci exhibiting linkage disequilibrium (LD); however, high LD can signal reduced recombination around genomic features such as chromosome inversions or sex-determining regions. Chromosome inversions and sex-determining regions are often involved in adaptation, allowing for the inheritance of co-adapted gene complexes and for the resolution of sexually antagonistic selection through sex-specific partitioning of genetic variants. Genomic features such as these can escape detection when loci with LD are removed; in addition, failing to account for these features can introduce bias to analyses. We examined patterns of LD using network analysis to identify an overlapping chromosome inversion and sex-determining region in chum salmon. The signal of the inversion was strong enough to show up as false population substructure when the entire dataset was analyzed, while the signal of the sex-determining region was only obvious after restricting genetic analysis to the sex chromosome. Understanding the extent and geographic distribution of inversions is now a critically important part of genetic analyses of natural populations. The results of this study highlight the importance of analyzing and understanding patterns of LD in genomic dataset and the perils of ignoring or excluding loci exhibiting LD.

## Introduction

The vast amount of genomic data now available allows research to move beyond analysis of allele frequency distributions among populations and into the effects of genomic structure and organization on adaptation and population structure. Genomic research has illuminated a range of evolutionary processes influencing adaptation. Single gene differences have been shown to have an important role in life-history variation: see for example the *GREB1L* gene that influences the timing of migration return in Chinook salmon (*Oncorhynchus tshawytscha*) and steelhead (*O. mykiss*) (Hess et al. 2016, Prince et al. 2017) and the *VGLL3* gene that maintains variation in age at maturity in Atlantic salmon (*Salmo salar*) (Barson et al. 2015). On a larger genomic scale, the importance of islands of divergence (e.g. Larson et al. 2017) and chromosome inversions (Wellenreuther and Bernatchez 2018) is increasingly recognized in non-model organisms, particularly in the context of sexually antagonistic selection (Kirkpatrick and Guerrero 2014, Barson et al. 2015). While islands of divergence can exhibit elevated linkage disequilibrium (LD) due to genetic hitchhiking, other features of genomic architecture such as inversions or sex-determining regions inhibit recombination, resulting in elevated LD and the inheritance of haplotype blocks of linked markers. Historically, many studies explicitly excluded loci exhibiting LD under the assumption that these markers provide redundant information (e.g., Larson et al. 2014), but with the current recognition of their importance and ability to detect genomic architecture, that practice is becoming obsolete.

Chromosome inversions have been associated with widely divergent adaptive variation ranging from social behavior (Thomas et al. 2008), life-history variation (Miller et al. 2012), alternative reproductive strategies (Küpper et al. 2015), and adaptation to different environments (Jones et al. 2012). Inversions have been implicated in the formation and divergence of sex chromosomes (Lemaitre et al. 2009) and establishment of reproductive barriers leading to speciation (Noor et al. 2001). Inversions often exhibit elevated divergence due to inhibited recombination, which can manifest as population substructure in genetic analyses (Tian et al. 2008, Arostegui et al. 2019), and the extended LD exhibited by large inversions facilitates their detection in marker-dense next generation sequencing projects (Kemppainen et al. 2015). An understanding of the extent and geographic distribution of inversions is now a critically important part of genetic analyses of natural populations.

Sex chromosomes are another genomic feature that often exhibits reduced recombination and elevated LD, with important evolutionary implications as males and females often exhibit different phenotypes and experience different selective pressures (Badyaev and Hill 2000, Tamate and Maekawa 2006). Elevated LD in sex chromosomes can be due to the presence of inversions (Lemaitre et al. 2009) or as a result of sex-specific patterns of recombination (Kijas et al. 2018). Reduced recombination can facilitate divergence between sex chromosomes under sexually antagonistic selection (Charlesworth 2018). Sexually antagonistic selection has been noted in a number of cases such as ornamental traits in poeciliid fishes (Anna Lindholm and Felix Breden 2002), optimal age at maturity in Atlantic salmon (Barson et al. 2015), and genetic variation for fitness in red deer (Foerster et al. 2007). When causal variants are located on the sex chromosome, sexually antagonistic selection can be resolved through portioning of genetic variation between sexes (Roberts et al. 2009).

With increasingly dense genomic data, sex-associated loci are likely to be found even if the sex-chromosome has not been identified; but, when unaccounted for, these sex-associated markers can bias genomic analyses (e.g. Benestan et al. 2017). Traditionally, sex-associated markers have been identified through genome-wide association studies (GWAS) of individuals with known sex or by identifying characteristics of heterogametic sex loci, such as presence/absence or genotypic frequencies where half of the samples are heterozygous and the other half are a single homozygous genotype (e.g. Star et al. 2016, McKinney et al. 2019). An alternative approach is to search for regions of extended LD which are often associated with sex-determining regions.

Identification of genomic features should be a routine step in genomic analyses to avoid biases outlined above (Benestan et al. 2017, Arostegui et al. 2019). Genome assemblies allow direct visualization of LD patterns along the chromosome, but most species do not have assembled genomes. In this case, network analysis can be performed on patterns of LD to identify genomic features (Kemppainen et al. 2015). Network analysis is likely to be particularly informative when multiple genomic features overlap, manifesting as a single region of high LD.

Here, we describe the detection and genetic architecture of multiple genomic features in chum salmon (*Oncorhynchus keta*) from western Alaska. We used patterns of LD combined with network analysis to identify a chromosomal inversion in chum salmon that co-occurs with the sex-determining region. The inversion exhibits spatial variation in frequency throughout western Alaska and includes putatively adaptive genes associated with life-history variation in other salmonids.

## Materials and Methods

### SNP Discovery and RAD sequencing

SNP discovery was conducted on 6 collections of chum salmon (48 samples each) using RAD sequencing (Figure 1). These collections were distributed among four major regions: Norton Sound, Yukon River, Kuskokwim River, and Nushagak River (Table 1). DNA was obtained from Alaska Department of Fish and Game (ADFG); the majority of these samples were analyzed in previous studies based on <100 SNPs (Seeb et al. 2011, Decovich et al. 2012). Sequencing libraries were prepared with the *SbfI* enzyme using a modification of the Rapture protocol (Best-RAD, Ali et al. 2016). Samples were sequenced on a HiSeq 4000 with paired-end 150 bp reads; 96 samples were sequenced per lane. Two rounds of sequencing were conducted, with the volume of DNA for each individual adjusted in the second round of sequencing to reduce variation in sequence reads per individual (Prince et al. 2017). Sequence data were processed with *STACKS* v1.47 (Catchen et al. 2011) using default settings with the following exceptions: process_radtags (-c -r -q -t 140), ustacks (-r --model_type bounded --bound_low 0 -- bound_high 0.01), cstacks (-n 2). The catalog of variation for *STACKS* was created using six individuals from each collection.

**Table 1.** Number of samples initially sequenced and retained after quality filtering for RAD and GT-seq datasets. Collections used to evaluate accuracy of the putative sex-determining region are marked with an asterisk *.

| Region | Collection | Initial RAD Samples | Retained RAD Samples | Initial GT-seq Samples | Retained GT-seq Samples |
| --- | --- | --- | --- | --- | --- |
| Norton Sound | Eldorado River | 48 | 48 | 83 | 82 |
| Norton Sound | Fish River* | 48 | 39 | 82 | 77 |
| Norton Sound | Kwiniuk River* | 0 | 0 | 82 | 74 |
| Yukon River | Nulato River | 48 | 38 | 83 | 80 |
| Yukon River | Otter Creek | 48 | 48 | 99 | 92 |
| Yukon River | East Fork Andreafsky River | 0 | 0 | 83 | 83 |
| Kuskokwim River | Holokuk River | 48 | 48 | 83 | 79 |
| Kuksokwim River | Aniak River | 0 | 0 | 82 | 79 |
| Nushagak River | Kokwok River | 48 | 46 | 83 | 79 |
| Nushagak River | Mulchatna River* | 0 | 0 | 82 | 73 |
|  |  | 288 | 267 | 842 | 798 |

**Figure 1.**
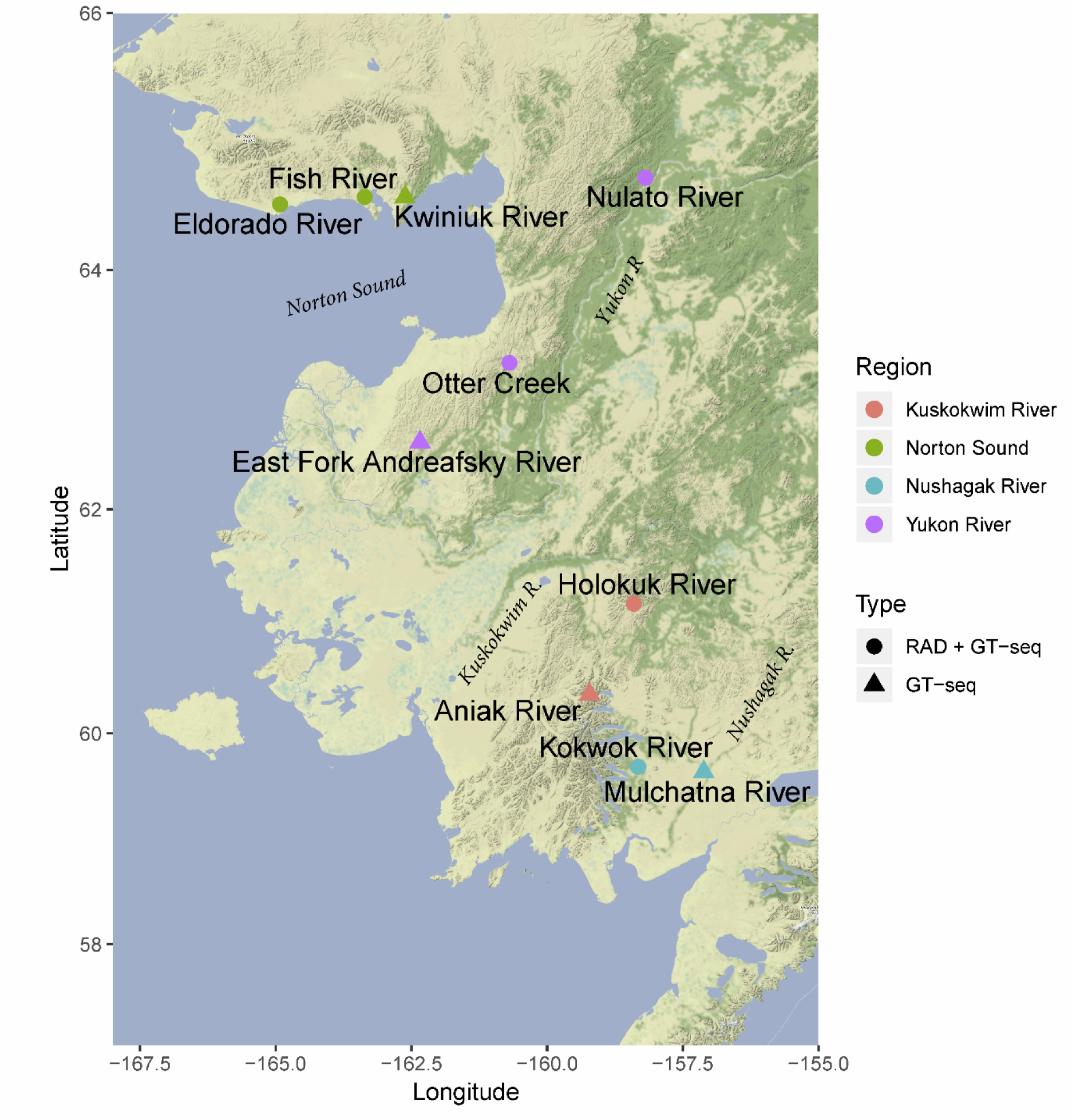
Map of sampling locations. Collections are colored by region, and shapes denote whether samples were genotyped using RAD and GT-seq or GT-seq only.

Loci and samples were filtered using an iterative process where poor-quality loci and samples were removed initially, the proportion of missing data was recalculated, and more stringent thresholds were applied for final filtering. A genotype rate threshold of 50% was initially applied to remove poor-quality loci. The genotype rate per sample was then estimated using the retained loci; samples were retained if they had a genotype rate of at least 75%. Allele frequencies and *F*_ST_ were estimated with *Genepop* (Rousset 2008) using the retained samples; loci with a minor allele frequency (MAF) of at least 0.05 were retained. *HDplot* (McKinney et al. 2017) was used to identify paralogs, which were excluded from further analysis because read depth was too low for accurate genotyping (McKinney et al. 2018). Singleton (non-paralogous) loci were retained for final analysis if they met a threshold of 90% genotype rate. Retained singleton loci were aligned against the rainbow trout *O. mykiss* genome (v_1.0, NCBI: GCA_002163495.1) using *bowtie2* (Langmead and Salzberg 2012) to allow investigation of genomic patterns of differentiation.

### Population Structure

Genomic features can manifest as population substructure when reduced recombination leads to fixation of alternate alleles over large genomic areas (e.g., Arostegui et al. 2019). Population structure was visualized using individual-based PCAs in R with Adegenet (Jombart 2008) to determine if any populations exhibited patterns of substructure. The populations included in this study spanned a large geographical region in Alaska, so population structure was examined within each region to prevent large-scale population structure from overwhelming any signal of genomic features. Populations exhibiting substructure were candidates for initial examination of LD.

### Identification of Genomic Structures

Genomic features were identified by examining LD between markers. Pairwise LD was estimated between SNPs using the r-squared method in *Plink* (v1.07, Purcell et al. 2007). Pairwise LD was plotted for each chromosome using R to identify regions of elevated LD. Network analysis and community detection were conducted in R using the igraph package (https://igraph.org/r) to identify groups of linked SNPs, hereafter referred to as ‘sets’. This was done to determine if multiple patterns of LD were present within a single genomic region. For network analysis, SNP pairs with an r^2^ < 0.3 were removed. SNPs that were in LD with fewer than 3 other SNPs were also excluded to reduce the number of SNPs that were linked only by close physical proximity. Following network analysis, chromosomal positions of SNPs in LD sets were examined to determine if they could be attributed to genomic structures. Finally, patterns of variation for SNPs within LD sets were visualized using PCA.

### High-throughput Assay Panel

Two putative, co-occurring genomic features were identified from the RADseq data (see results): a chromosome inversion and a sex-determining region.

GT-seq assays were developed for 39 loci diagnostic for the putative inversion, and 21 loci diagnostic for the putative sex-determining region. Multiple loci for each genomic feature were included to provide a control for genotyping error and because some loci were expected to be lost during panel optimization (described below). Filters were applied prior to and during primer design to remove loci that were likely to amplify off-target sequence with GT-seq following the methods of McKinney et al. (2019); this included identifying transposable elements and primers that align to multiple genomic regions. Loci with SNPs within 20 bp of the end of the RAD tag required genomic sequence past the RAD tag for primer design. We obtained a draft copy of the chum salmon genome from Ben Koop and aligned RAD tag sequences to the unassembled scaffolds using *bowtie2*; custom perl scripts were used to obtain flanking sequence for primer design. Primers were designed using *batch primer3* (You et al. 2008). Amplicons for retained primers were then examined to ensure that SNPs were contained within the first 100 bp of the amplicon to facilitate downstream single-end sequencing.

Two rounds of panel optimization were conducted to identify and remove loci that did not perform well. Each round of optimization was conducted using 48 individuals sequenced on a MiSeq with 150 bp paired-end sequence. DNA was extracted and sequencing libraries were prepared following the methods of Campbell et al. (2015). Sequencing data were processed and genotyped using *GTscore* (https://github.com/gjmckinney/GTscore, McKinney et al. 2019). After each round of sequencing, the number of reads amplified by each primer was counted, as well as the proportion of reads that contained both the primer and bioinformatics probes for a locus. Loci with excessive amplification and off-target amplification were removed following the methods of McKinney et al. (2019).

An expanded sample of individuals was genotyped using the GT-seq panel to better characterize genomic structure across the region. This included additional individuals from the RAD ascertainment collections as well as additional collections from Norton Sound, and the Yukon, Kuskokwim, and Nushagak rivers (Table 1). Populations and individuals with paired sex data were preferentially chosen to assess concordance between phenotypic sex and the putative sex-determining region. Sequencing libraries were prepared as above. A total of 871 samples (842 plus 39 sequenced twice with GT-seq for quality control) were sequenced on a single lane of a HiSeq 4000 with 100 bp single-end sequencing. These samples were also genotyped for an additional 478 markers on this sequencing lane as part of a genetic stock identification (GSI) project (McKinney et al., unpubl.). All GT-seq loci were used for evaluating sample quality even though loci developed for the GSI project are not included in this study. Samples were evaluated for quality based on a minimum 90% genotype rate and visualization of allele scatter plots.

## Results

### SNP Discovery and RAD sequencing

*STACKS* initially outputted 222,668 SNPs within 94,002 RAD tags. A total of 30,006 SNPs within 22,693 RAD tags were retained after applying all filters (Table S1). Of these loci, 13,015 SNPs within 10,821 RAD tags were aligned to the rainbow trout genome. After all filtering steps, 267 individuals were retained for SNP discovery.

### Identification of Genomic Structures

Clear patterns of population substructure were apparent in the Yukon River from individual-based PCAs on all 13,015 markers (Figure 2). No substructure was visible in individual PCAs for other regions (Figure S1). The loci driving population substructure within Yukon River collections were part of a region of elevated LD on Omy28 (Figure 3A); this corresponds to linkage group 29 on the chum salmon linkage map (Waples et al. 2016). Network analysis identified three distinct sets of loci that exhibited high LD (Figure 3B). Loci in set 1 spanned from 437 Kb to 20.3 Mb and the loci in set 2 spanned from 1.6 Mb to 21.3 Mb. The loci in set 3 spanned only a 2 Kb region, suggesting that their high LD is likely due only to close physical proximity. Therefore, these loci were excluded from further analysis.

**Figure 2.**
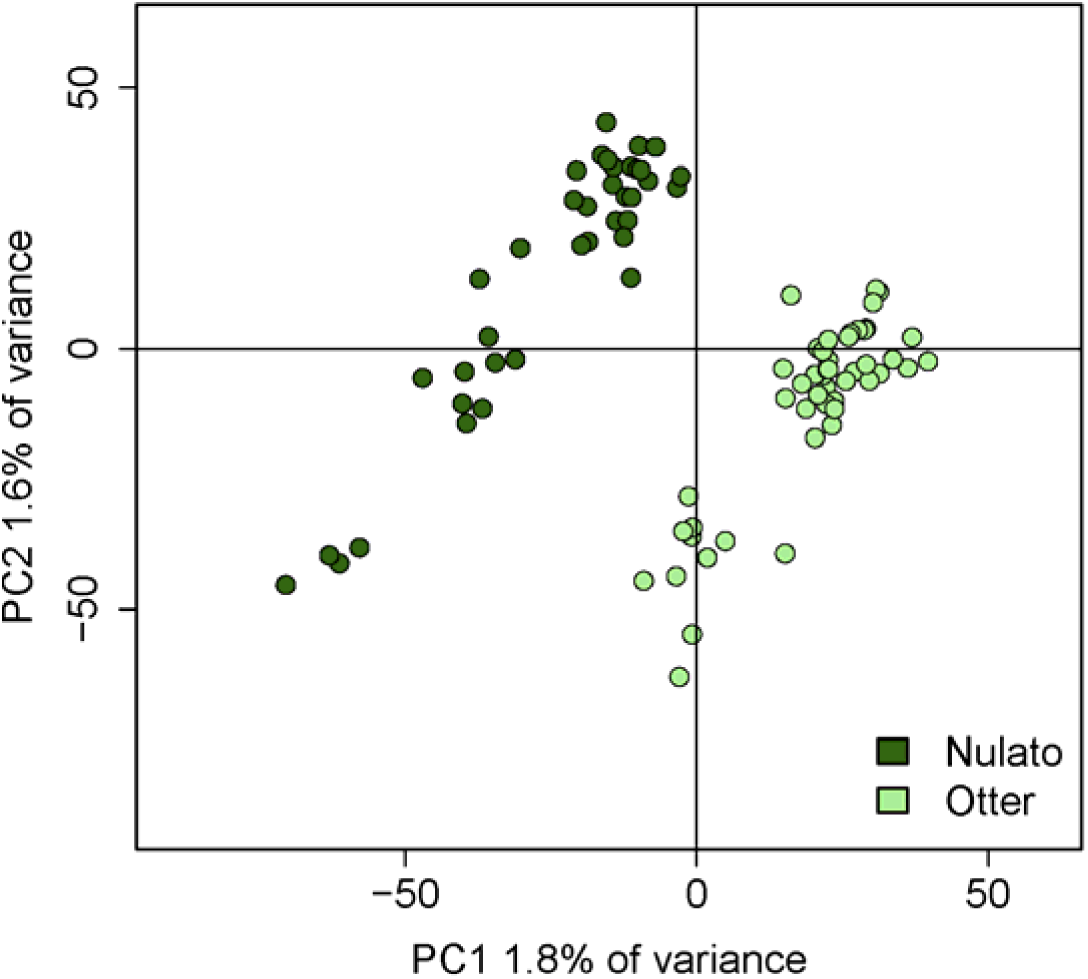
Individual PCA using all loci for Yukon River collections. Both collections exhibit patterns of substructure.

**Figure 3.**
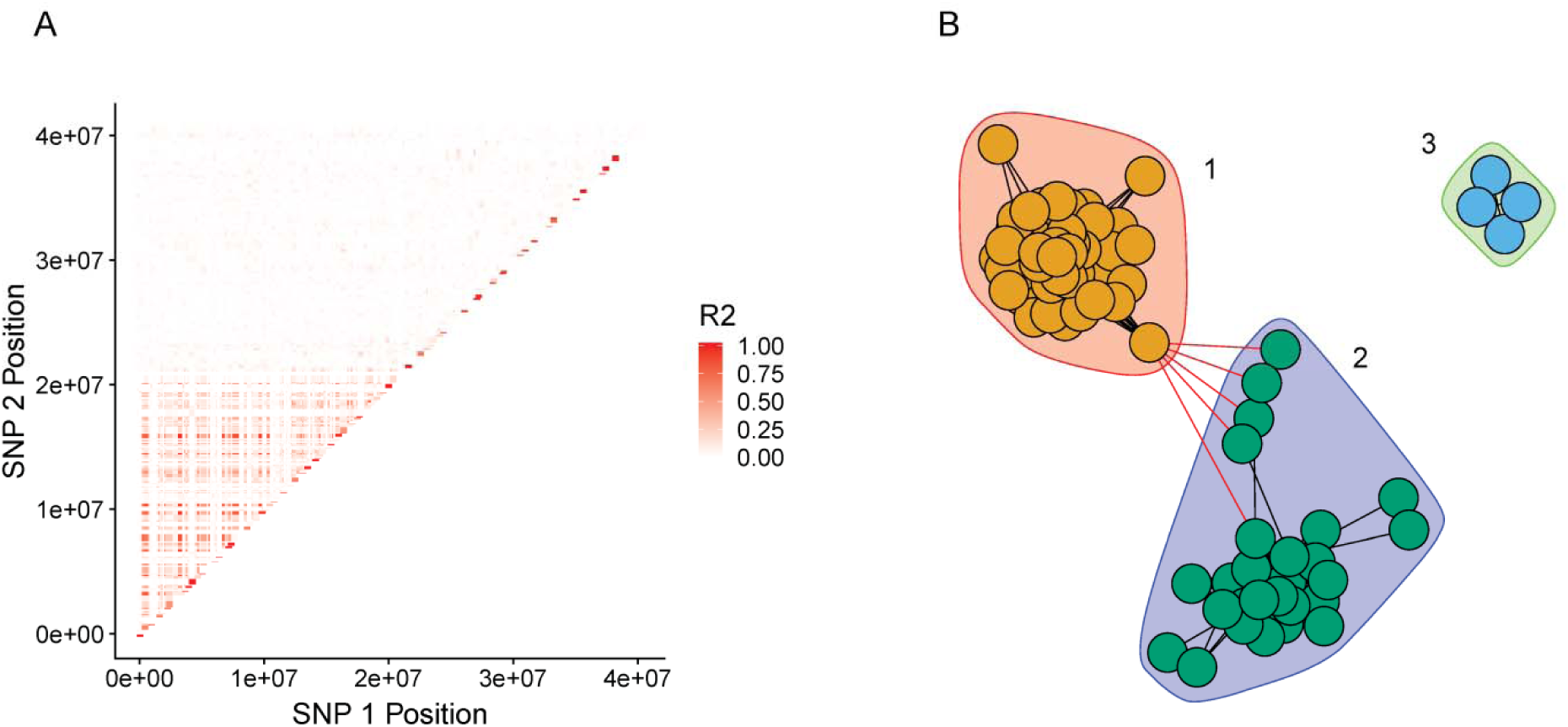
A) Plot of linkage disequilibrium (pairwise r^2^) on *O. mykiss* chromosome Omy28. Each point is a SNP pair colored by LD. The pattern of elevated LD spans 20 Mb of chromosome 28. B) Network analysis with community detection identified three distinct sets of loci contributing to LD on Omy28. Set 1 (red background) has 51 loci, set 2 (purple background) has 27 loci, and set 3 (green background) has 4 loci. Loci in sets 1 and 2 span the entire 20 Mb while loci in set 3 are linked due to close physical proximity.

We then plotted all RADseq samples for all loci from Omy28. The PCA of Omy28 revealed clustering of samples that was driven by the loci in sets 1 and 2; PCA loadings show that Axis 1 was primarily driven by loci in set 1, and axis 2 was primarily driven by loci in set 2 (Figure 4A).

**Figure 4.**
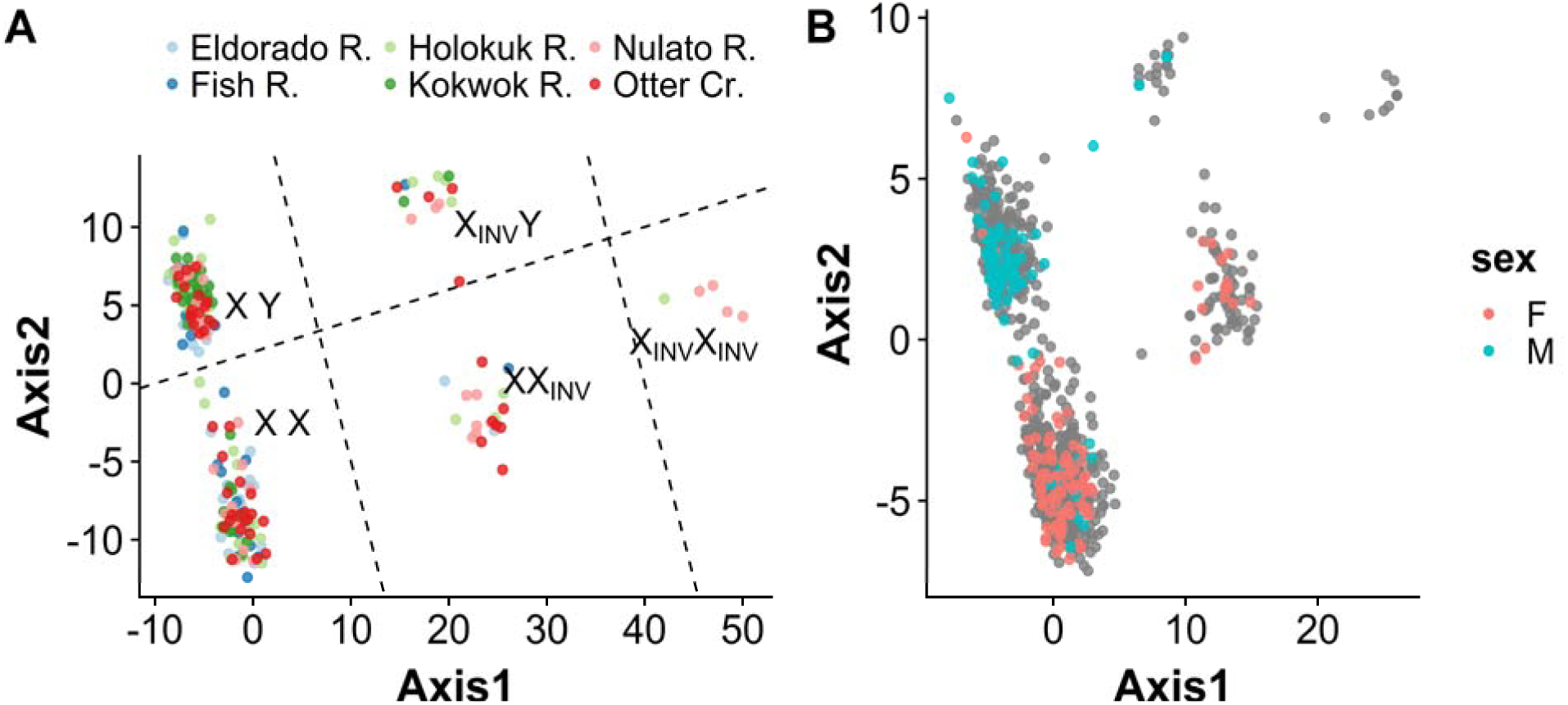
A) Individual PCA of RADseq samples using all loci from Omy28. Examination of locus loadings show that Axis 1 is primarily driven by loci in set 1 (inversion loci) and axis 2 is primarily driven by loci in set 2 (sex-associated loci). Labels for each cluster of individuals denote the putative chromosome type of individuals within the cluster with respect to inversion and sex. B) Individual PCA of RADseq and GT-seq samples using loci successfully developed into GT-seq assays. Samples are color coded by phenotypic sex.

Genotype patterns within a genomic feature can give clues to the type of genomic feature. Genotypes for loci in sets 1 and 2 (including those genotyped with GT-seq; see below) were plotted by SNP set and position to better visualize genotype patterns (Figure 5); this plot revealed patterns consistent with multiple genomic features within this region. Loci in set 1 exhibited three genotype classes (both homozygous and heterozygous) and near-complete linkage for loci spanning 20 Mb, which is consistent with a genomic inversion that is variable within populations (e.g. Arostegui et al. 2019). We assume that the inversion was the least common form but this is not always the case (Arostegui et al. 2019). This inversion was present in all collections, but its frequency varied by region, with Yukon and Kuskokwim rivers having the highest frequency of the inversion (Table 2).

**Table 2.** Frequency of the chromosome inversion and sex-assignment accuracy by collection. Samples were assigned an inversion type (+/-) and sex based on PCA clustering. For collections with phenotypic sex data, phenotypes were compared to sex assigned through clustering to assess accuracy.

| Region | Collection | Data Source | ++ | +- | -- | Freq (-) | M cluster | F cluster | Sex Assignment Accuracy |
| --- | --- | --- | --- | --- | --- | --- | --- | --- | --- |
| Norton Sound | Eldorado River | RAD/GT-seq | 96% | 4% | 0% | 2% | 50 | 80 | NA |
| Norton Sound | Fish River | RAD/GT-seq | 90% | 10% | 0% | 6% | 54 | 62 | 84% |
| Norton Sound | Kwiniuk River | GT-seq | 92% | 8% | 0% | 4% | 30 | 44 | 89% |
| Yukon R. | Nulato River | RAD/GT-seq | 73% | 23% | 4% | 22% | 65 | 53 | NA |
| Yukon R. | Otter Creek | RAD/GT-seq | 76% | 22% | 1% | 16% | 65 | 75 | NA |
| Yukon R. | East Fork Andreafsky | GT-seq | 83% | 17% | 0% | 10% | 37 | 46 | NA |
| Kuskokwim R. | Holokuk River | RAD/GT-seq | 89% | 10% | 1% | 7% | 54 | 73 | NA |
| Kuskokwim R. | Aniak River | GT-seq | 85% | 15% | 0% | 9% | 41 | 38 | NA |
| Nushagak R. | Kokwok River | RAD/GT-seq | 96% | 4% | 0% | 2% | 96 | 29 | NA |
| Nushagak R. | Mulchatna River | GT-seq | 90% | 10% | 0% | 5% | 37 | 36 | 99% |

**Figure 5.**
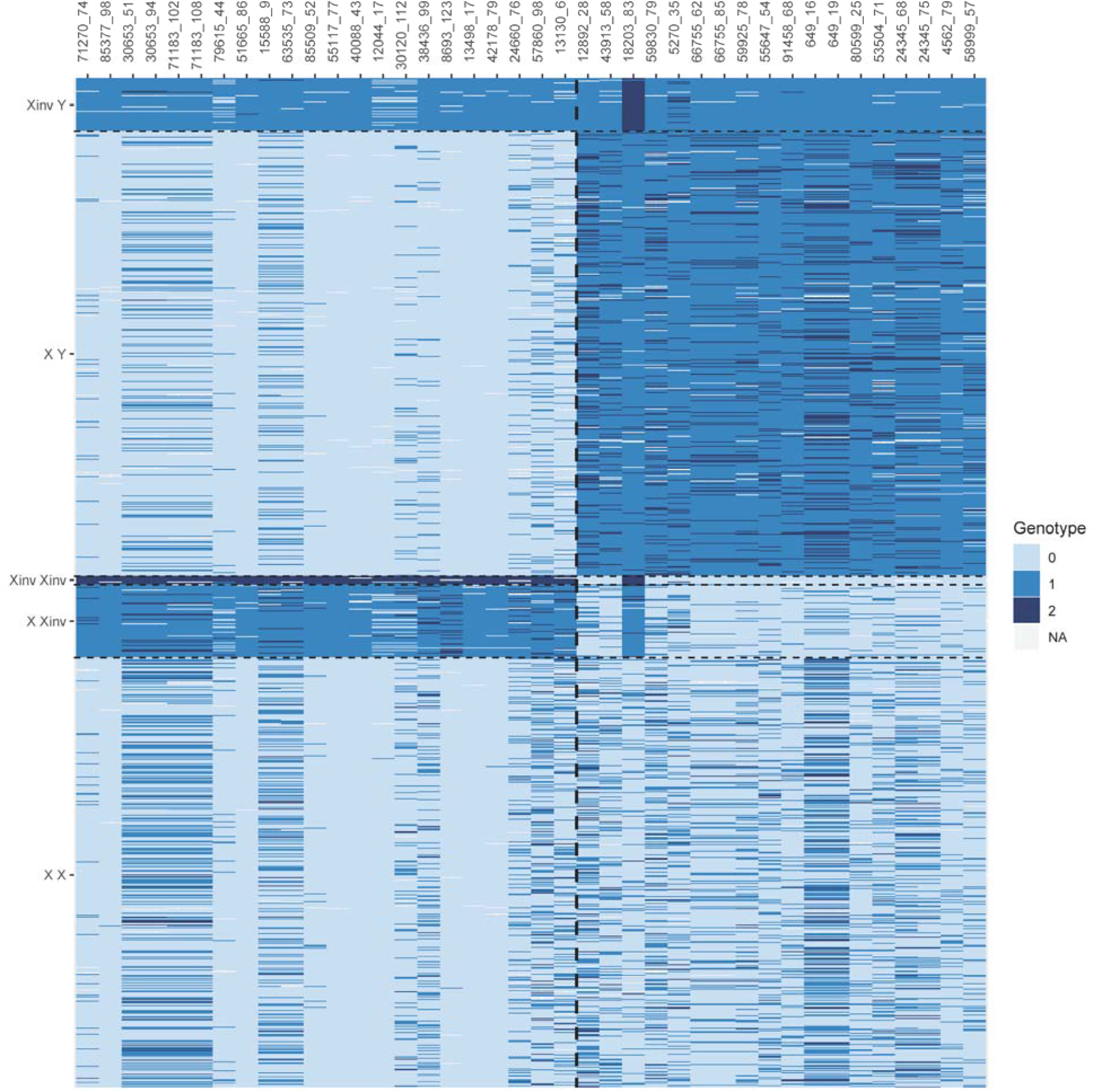
Genotypes for combined RADseq and GT-seq samples. Samples are in rows ordered by sex (inferred from PCA cluster) and inversion type and loci are in columns. Loci were separated by genomic feature to aid in visualization: inversion-associated loci are to the left of the dashed dividing line and sex-associated loci to the right. Within each genomic feature, loci are ordered by position. Individual genotypes are color coded with 0 and 2 representing alternate homozygous genotypes and 1 being a heterozygous genotype. Prefix of Oke_uwRAD was dropped from marker names for brevity.

Loci in set 2 exhibited a more complex pattern that is consistent with a sex-determining region. Putative males in the top half of Figure 5 and loci to the right of the dividing line show a pattern of very high heterozygosity with homozygous genotypes almost entirely of one class rather than both homozygous genotypes being equally represented. Putative females in the bottom half of Figure 5 with loci to the right of dividing line show a pattern of high homozygosity for the alternate allele found in males, and all three genotype classes are present. This overall pattern is consistent with fixation of one allele on the Y-chromosome with the alternate allele at high frequency, but not fixed, on the ancestral X-chromosome. These patterns are consistent with an XY sex-determining region and balanced sex ratios. This finding was further supported by phenotypic sex data that were available for 37 individuals from Fish River; 13 of 15 phenotypic females grouped with the putative females (based on set 2 loci) and 18 of 22 phenotypic males grouped with putative males (Table 2, Figure 4B). Note that Figure 4B includes all available RADseq and GT-seq samples.

### High-throughput Panel for Screening Genomic Structures

Development of the GT-seq panel involved filtering steps during primer design and panel optimization. After evaluation and optimization, the final GT-seq panel contained 22 SNPs diagnostic for a putative inversion and 18 SNPs diagnostic for a putative sex-determining region. Primer sequences for final loci are located in Supplemental file S1. A total of 43 GT-seq samples were removed for failing quality control; 19 samples had <90% genotype rate and an additional eight had broad allelic scatter due to reduced read depth (Figure S2B), five samples showed elevated heterozygosity and indistinct allelic clusters suggesting contamination (Figure S2C), one sample showed four clusters consistent with triploidy (Figure S2D), two samples showed five clusters consistent with tetraploidy (Figure S2E), and one sample had no heterozygous genotypes and was likely a different species of salmon (Figure S2F).

Inversion type and sex were assigned by clustering samples with PCA (Figure 4B); this PCA mirrored the results from RADseq analysis shown in Figure 4A. Results for inversion and sex are reported for the full dataset with the RADseq and GT-seq samples combined. Phenotypic sex data were available for samples in five of the collections examined; however, systemic errors in phenotypic records were identified for the East Fork Andreafsky and Aniak Rivers populations, causing phenotypes for these populations to be removed from analysis. The average concordance between phenotypic sex and cluster sex for the remaining combined RADseq and GT-seq datasets was 90% (range 84%-99%, Table 2).

Similar to the RAD data alone, within the full dataset the inversion was found in every collection but showed regional variation in frequency. Inversion frequency ranged from 2% to 22% and was most common in the Lower Yukon collections followed by the Kuskokwim River collections (Table 2). Outside of the Yukon and Kuskokwim rivers the inversion reached a maximum frequency of only 6% (Table 2).

## Discussion

Genomic structures including inversions and sex-determining regions are increasingly recognized to contribute to adaptive variation and population structure (Benestan et al. 2017, Wellenreuther and Bernatchez 2018). Large inversions are associated with adaptation and life-history variation across taxa, including ecotypes in mosquitos (*Anopheles* sp.) (Ayala et al. 2017), migration vs. residency in rainbow trout (*Oncorhynchus mykiss*) (Pearse et al. 2018), and annual vs. perennial yellow monkey flowers (*Mimulus guttatus*) (Lowry and Willis 2010). Inhibited recombination within inverted regions increases divergence between chromosomal forms and can drive patterns of population structure (Tian et al. 2008, Arostegui et al. 2019). A similar pattern has been observed for sex chromosomes where sex-associated markers can drive patterns of population structure (Benestan et al. 2017). We identified two genomic structures in chum salmon - a chromosome inversion and sex-determining region - that occur in the same genomic region. The signal of the chromosome inversion was strong enough to cause population structuring when all markers were used for PCA; however, population structure due to the sex-determining region was only visible in the chromosome-specific PCA.

### Identification of Genomic Structures Using LD

Population substructure in the RADseq data shaped by the inversion and sex-determining region was only understood through a combination of LD and network analysis. Both the inversion and sex-determining region exhibited LD spanning the same genomic region, which would be difficult to tease out manually. Network analysis and community detection was a simple method to automate detection of different groups of linked markers that contributed to this overall pattern of LD. Previous work has shown that a combination of LD and network analyses facilitates detection of genomic features even in the absence of reference genome (Kemppainen et al. 2015). Here we demonstrated that LD and network analysis can be used to tease apart multiple genomic features within the same chromosomal region.

The extended LD combined with the specific genotype patterns observed for high-LD loci suggests a genomic inversion that is co-occurring with the sex-determining region in chum salmon. The genotype patterns also suggest that the inversion arose on the X chromosome. An inversion arising on the X chromosome should result in 5 different chromosomal combinations: XX, XX_INV_, X_INV_X_INV_, XY, X_INV_Y. This agrees with the observed pattern of 5 sample clusters on the individual PCA (Figure 4). If the inversion had arisen on the Y chromosome only three clusters would be expected: XX, XY, XY_INV_. Levels of LD for sex-associated loci varied depending on the X-chromosome type, suggesting different levels of divergence between the Y-chromosome and the ancestral and inverted X-chromosomes.

### Chromosome Inversion

A genomic inversion on the X chromosome containing the sex-determining region has interesting evolutionary implications. Females can exhibit all three genotypes of the inversion, while males can be homozygous for the ancestral form or heterozygous for the inversion but never homozygous for the inversion. Genomic inversions can lead to fixation of genetic variants on alternate chromosome forms and facilitate adaptive divergence (reviewed in Wellenreuther and Bernatchez 2018). In salmonids, an inversion on Omy05 is associated with variation in migrant vs. resident life history in rainbow trout in both saltwater (Miller et al. 2012, Pearse et al. 2014, Pearse et al. 2018) and freshwater systems (Arostegui et al. 2019). We cannot determine if the inversion we identified in chum salmon is adaptive with the data currently available but one intriguing result is that the inversion contains the Greb1L gene. This gene has been associated with variation in migration timing in Chinook salmon and rainbow trout (Hess et al. 2016, Prince et al. 2017, Micheletti et al. 2018). Chum salmon exhibit summer and fall life histories that differ in migration timing; however, this study examined only summer run chum salmon. Further study of individual migration timing could determine whether there is an association between inversion type and return date in chum salmon. We also recommend follow up work with whole-genome sequencing combined with individual metadata to better determine if this inversion is adaptive, and what genes and gene variants may be involved (i.e. Pearse et al. 2018).

### Sex-Determining Region

Collections exhibited variation in concordance between phenotypic and inferred sex. There are two possible sources of error that may explain this: error in phenotypic sex assignment and error in sample records. The populations in this study were sexed visually which can be unreliable, particularly if fish were caught before secondary sex characteristics were fully developed (Lozorie and McIntosh 2014). Despite this, overall concordance between phenotypic sex and sex determined by PCA cluster was high (90% overall and 99% in the Mulchatna collection) suggesting that this is the true sex determining region. Phenotypes for two populations in this study, East Fork Andreafsky and Aniak, were excluded due to record errors. Future studies of this genomic region should involve new collections of samples with careful attention to the accurate pairing of phenotypic data and tissue samples.

## Conclusion

Examining genome-wide patterns of LD is an important tool for evolutionary analysis. Many studies explicitly exclude loci with LD; these may be missing important patterns of genomic variation. We identified a co-locating inversion and sex-determining region in chum salmon by performing network analysis on patterns of LD. The signal of these features was strong enough to drive PCA of the full dataset, resulting in false population structure. Attempting to identify and account for genomic features should be standard practice in genome-scale datasets.

## Acknowledgements

We would like to thank Bill Templin, Chris Habicht, and Sara Gilk-Baumer from the Alaska Department of Fish and Game (ADFG) for their support and assistance during this project. We would also like to thank the many biologists at ADFG who collected the samples used in this project. Funding for this project was provided by the Pollock Cooperative Conservation and Research Center and the Bureau of Energy Management (CFDA 15.668; award # F12AF00424). The statements, findings, and conclusions are the authors’ and do not necessarily reflect the views of the funding entities.

## Supplemental

Supplemental File S1: Primers for GT-seq loci developed in this project.

**Table S1.**
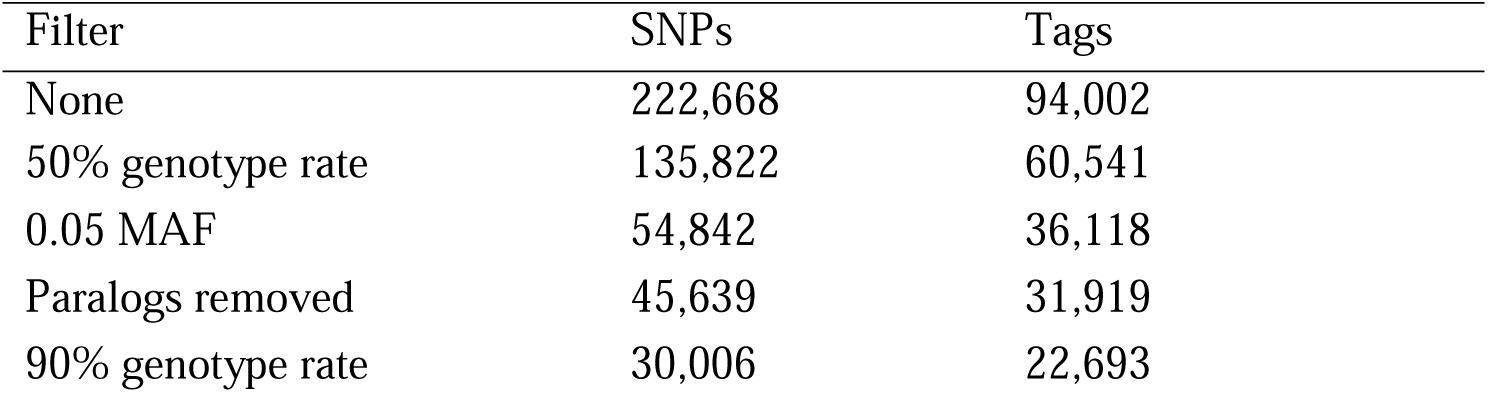
Results of RAD marker filtering by step. The first column lists the filter applied. The SNPs column lists the number of SNPs retained after this filter while the Tags column lists the number of RAD tags retained after this filter.

**Figure S1.**
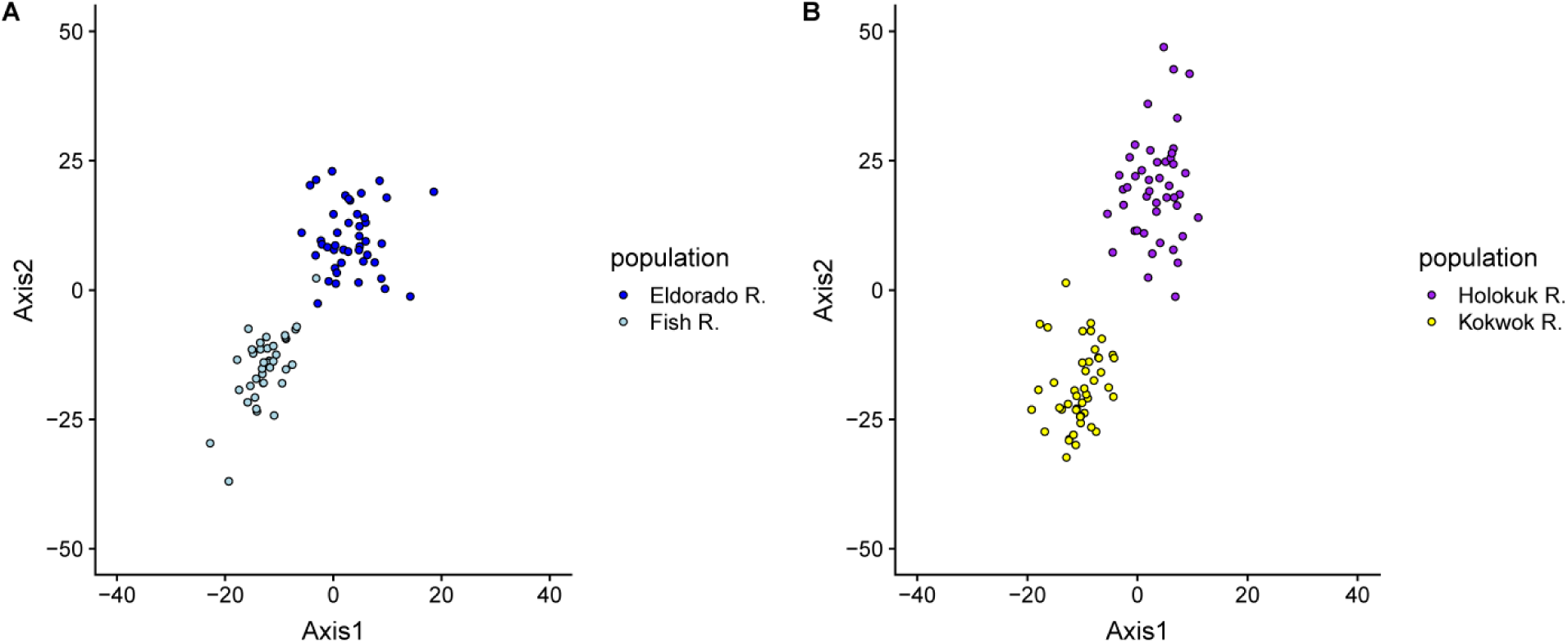
Individual PCAs using all loci for A) Norton Sound collections and B) Kuskokwim and Nushagak collections. Among-population divergence is apparent but no populations exhibit patterns of substructure.

**Figure S2.**
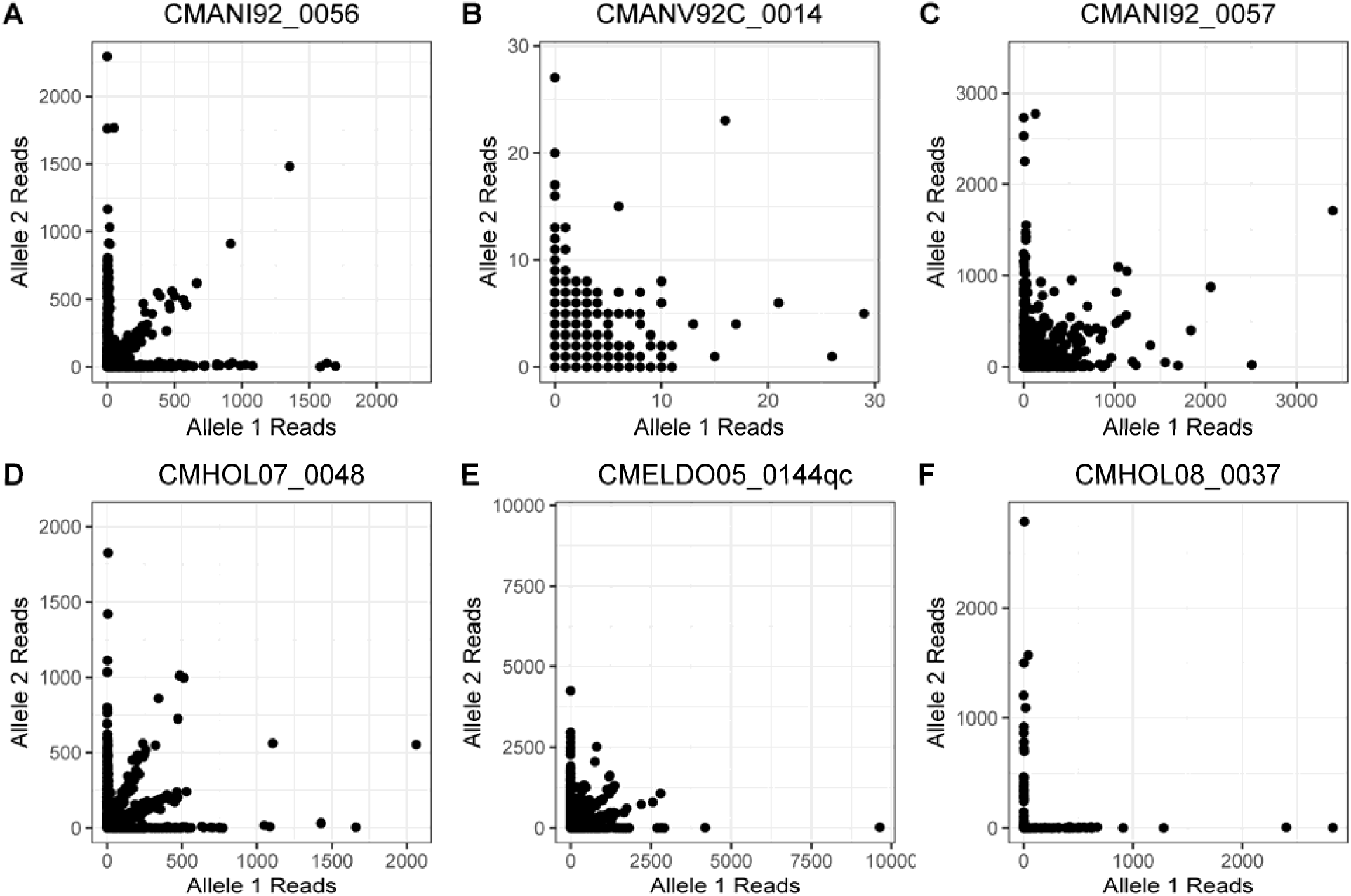
Example scatter plots of allele reads for samples. The expected pattern is shown in A Genotypes for distinct clusters with homozyous genotypes primarily have reads for a single allele, reads for the alternate allele are presumably due to sequencing error, and heterozygous genotypes have approximately equal reads for each allele. The scatterplot in B is indistinct due to very low read depth. The scatterplot in C fails to form distinct clusters despite high read depth, suggesting sample contamination. The scatterplot in D has two clusters of heterozygous genotypes following 1:2 and 2:1 ratios consistent with a triploid individual. The scatterplot in E has three clusters of heterozygous genotypes at 1:3, 2:2, and 3:1 ratios consistent with a tetraploid individual. The scatterplot in F has no heterozygous genotypes which is consistent with a sample from another species.

## References

Ali, O. A., S. M. O’Rourke, S. J. Amish, M. H. Meek, G. Luikart, C. Jeffres, and M. R. Miller. 2016. RAD capture (Rapture): flexible and efficient sequence-based genotyping. Genetics 202:389–400.

Anna Lindholm, and Felix Breden. 2002. Sex Chromosomes and Sexual Selection in Poeciliid Fishes. The American Naturalist 160:S214–S224.

Arostegui, M. C., T. P. Quinn, L. W. Seeb, J. E. Seeb, and G. J. McKinney. 2019. Retention of a chromosomal inversion from an anadromous ancestor provides the genetic basis for alternative freshwater ecotypes in rainbow trout. Mol Ecol.

Ayala, D., P. Acevedo, M. Pombi, I. Dia, D. Boccolini, C. Costantini, F. Simard, and D. Fontenille. 2017. Chromosome inversions and ecological plasticity in the main African malaria mosquitoes. Evolution; international journal of organic evolution 71:686–701.

Badyaev, A. V., and G. E. Hill. 2000. THE EVOLUTION OF SEXUAL DIMORPHISM IN THE HOUSE FINCH. I. POPULATION DIVERGENCE IN MORPHOLOGICAL COVARIANCE STRUCTURE. Evolution 54:1784–1794.

Barson, N. J., T. Aykanat, K. Hindar, M. Baranski, G. H. Bolstad, P. Fiske, C. Jacq, A. J. Jensen, S. E. Johnston, S. Karlsson, M. Kent, T. Moen, E. Niemela, T. Nome, T. F. Naesje, P. Orell, A. Romakkaniemi, H. Saegrov, K. Urdal, J. Erkinaro, S. Lien, and C. R. Primmer. 2015. Sex-dependent dominance at a single locus maintains variation in age at maturity in salmon. Nature 528:405–408.

Benestan, L., J. S. Moore, B. J. G. Sutherland, J. Le Luyer, H. Maaroufi, C. Rougeux, E. Normandeau, N. Rycroft, J. Atema, L. N. Harris, R. F. Tallman, S. J. Greenwood, F. K. Clark, and L. Bernatchez. 2017. Sex matters in massive parallel sequencing: Evidence for biases in genetic parameter estimation and investigation of sex determination systems. Mol Ecol 26:6767–6783.

Campbell, N. R., S. A. Harmon, and S. R. Narum. 2015. Genotyping-in-Thousands by sequencing (GT-seq): a cost effective SNP genotyping method based on custom amplicon sequencing. Mol Ecol Resour 15:855–867.

Catchen, J. M., A. Amores, P. Hohenlohe, W. Cresko, and J. H. Postlethwait. 2011. Stacks: building and genotyping loci de novo from short-read sequences. G3: Genes, Genomes, Genetics 1:171–182.

Charlesworth, D. 2018. The Guppy Sex Chromosome System and the Sexually Antagonistic Polymorphism Hypothesis for Y Chromosome Recombination Suppression. Genes 9:264.

Decovich, N. A., T. H. Dann, S. D. R. Olive, H. L. Liller, E. K. C. Fox, J. R. Jasper, E. L. Chenoweth, C. Habicht, and W. D. Templin. 2012. Chum Salmon Baseline for the Western Alaska Salmon Stock Identification Program. Alaska Department of Fish and Game, Division of Commercial Fisheries Alaska Department of Fish and Game, Special Publication No. 12-26, 110p.

Foerster, K., T. Coulson, B. C. Sheldon, J. M. Pemberton, T. H. Clutton-Brock, and L. E. B. Kruuk. 2007. Sexually antagonistic genetic variation for fitness in red deer. Nature 447:1107.

Hess, J. E., J. S. Zendt, A. R. Matala, and S. R. Narum. 2016. Genetic basis of adult migration timing in anadromous steelhead discovered through multivariate association testing. Proc Biol Sci 283.

Jombart, T. 2008. adegenet: a R package for the multivariate analysis of genetic markers. Bioinformatics 24:1403–1405.

Jones, F. C., M. G. Grabherr, Y. F. Chan, P. Russell, E. Mauceli, J. Johnson, R. Swofford, M. Pirun, M. C. Zody, S. White, E. Birney, S. Searle, J. Schmutz, J. Grimwood, M. C. Dickson, R. M. Myers, C. T. Miller, B. R. Summers, A. K. Knecht, S. D. Brady, H. Zhang, A. A. Pollen, T. Howes, C. Amemiya, P. Broad Institute Genome Sequencing, T. Whole Genome Assembly, J. Baldwin, T. Bloom, D. B. Jaffe, R. Nicol, J. Wilkinson, E. S. Lander, F. Di Palma, K. Lindblad-Toh, and D. M. Kingsley. 2012. The genomic basis of adaptive evolution in threespine sticklebacks. Nature 484:55.

Kemppainen, P., C. G. Knight, D. K. Sarma, T. Hlaing, A. Prakash, Y. N. Maung Maung, P. Somboon, J. Mahanta, and C. Walton. 2015. Linkage disequilibrium network analysis (LDna) gives a global view of chromosomal inversions, local adaptation and geographic structure. Mol Ecol Resour 15:1031–1045.

Kijas, J., S. McWilliam, M. Naval Sanchez, P. Kube, H. King, B. Evans, T. Nome, S. Lien, and K. Verbyla. 2018. Evolution of Sex Determination Loci in Atlantic Salmon. Scientific Reports 8:5664.

Kirkpatrick, M., and R. F. Guerrero. 2014. Signatures of Sex-Antagonistic Selection on Recombining Sex Chromosomes. Genetics 197:531–541.

Koop, B. F. In Prep. Chum salmon genome.

Küpper, C., M. Stocks, J. E. Risse, N. dos Remedios, L. L. Farrell, S. B. McRae, T. C. Morgan, N. Karlionova, P. Pinchuk, Y. I. Verkuil, A. S. Kitaysky, J. C. Wingfield, T. Piersma, K. Zeng, J. Slate, M. Blaxter, D. B. Lank, and T. Burke. 2015. A supergene determines highly divergent male reproductive morphs in the ruff. Nature Genetics 48:79.

Langmead, B., and S. L. Salzberg. 2012. Fast gapped-read alignment with Bowtie 2. Nat Methods 9:357–359.

Larson, W. A., M. T. Limborg, G. J. McKinney, D. E. Schindler, J. E. Seeb, and L. W. Seeb. 2017. Genomic islands of divergence linked to ecotypic variation in sockeye salmon. Mol Ecol 26:554–570.

Larson, W. A., L. W. Seeb, M. V. Everett, R. K. Waples, W. D. Templin, and J. E. Seeb. 2014. Genotyping by sequencing resolves shallow population structure to inform conservation of Chinook salmon (Oncorhynchus tshawytscha). Evol Appl 7:355–369.

Lemaitre, C., M. D. V. Braga, C. Gautier, M.-F. Sagot, E. Tannier, and G. A. B. Marais. 2009. Footprints of inversions at present and past pseudoautosomal boundaries in human sex chromosomes. Genome Biology and Evolution 1:56–66.

Lowry, D. B., and J. H. Willis. 2010. A Widespread Chromosomal Inversion Polymorphism Contributes to a Major Life-History Transition, Local Adaptation, and Reproductive Isolation. PLOS Biology 8:e1000500.

Lozorie, J. D., and B. C. McIntosh. 2014. Sonar esimation of salmon passage in the Yukon River near Pilot Station, 2012. Alaska Department of Fish and Game.

McKinney, G. J., C. E. Pascal, W. D. Templine, S. E. Gilk-Baumer, T. H. Dann, L. W. Seeb, and J. E. Seeb. 2019. Dense SNP panels resolve closely related Chinook salmon populations. Canadian Journal of Fisheries and Aquatic Sciences Accepted.

McKinney, G. J., R. K. Waples, C. E. Pascal, L. W. Seeb, and J. E. Seeb. 2018. Resolving allele dosage in duplicated loci using genotyping-by-sequencing data: A path forward for population genetic analysis. Mol Ecol Resour 18:570–579.

McKinney, G. J., R. K. Waples, L. W. Seeb, and J. E. Seeb. 2017. Paralogs are revealed by proportion of heterozygotes and deviations in read ratios in genotyping-by-sequencing data from natural populations. Mol Ecol Resour 17:656–669.

Micheletti, S. J., J. E. Hess, J. S. Zendt, and S. R. Narum. 2018. Selection at a genomic region of major effect is responsible for evolution of complex life histories in anadromous steelhead. Bmc Evolutionary Biology 18:140.

Miller, M. R., J. P. Brunelli, P. A. Wheeler, S. Liu, C. E. Rexroad, 3rd, Y. Palti, C. Q. Doe, and G. H. Thorgaard. 2012. A conserved haplotype controls parallel adaptation in geographically distant salmonid populations. Mol Ecol 21:237–249.

Noor, M. A. F., K. L. Grams, L. A. Bertucci, and J. Reiland. 2001. Chromosomal inversions and the reproductive isolation of species. Proceedings of the National Academy of Sciences 98:12084–12088.

Pearse, D. E., N. J. Barson, T. Nome, G. Gao, M. A. Campbell, A. Abadía-Cardoso, E. C. Anderson, D. E. Rundio, T. H. Williams, K. A. Naish, T. Moen, S. Liu, M. Kent, D. R. Minkley, E. B. Rondeau, M. S. O. Brieuc, S. Rød Sandve, M. R. Miller, L. Cedillo, K. Baruch, A. G. Hernandez, G. Ben-Zvi, D. Shem-Tov, O. Barad, K. Kuzishchin, J. C. Garza, S. T. Lindley, B. F. Koop, G. H. Thorgaard, Y. Palti, and S. Lien. 2018. Sex-dependent dominance maintains migration supergene in rainbow trout. bioRxiv:504621.

Pearse, D. E., M. R. Miller, A. Abadia-Cardoso, and J. C. Garza. 2014. Rapid parallel evolution of standing variation in a single, complex, genomic region is associated with life history in steelhead/rainbow trout. Proc Biol Sci 281:20140012.

Prince, D. J., S. M. O’Rourke, T. Q. Thompson, O. A. Ali, H. S. Lyman, I. K. Saglam, T. J. Hotaling, A. P. Spidle, and M. R. Miller. 2017. The evolutionary basis of premature migration in Pacific salmon highlights the utility of genomics for informing conservation. Sci Adv 3:e1603198.

Purcell, S., B. Neale, K. Todd-Brown, L. Thomas, M. A. Ferreira, D. Bender, J. Maller, P. Sklar, P. I. de Bakker, M. J. Daly, and P. C. Sham. 2007. PLINK: a tool set for whole-genome association and population-based linkage analyses. Am J Hum Genet 81:559–575.

Roberts, R. B., J. R. Ser, and T. D. Kocher. 2009. Sexual Conflict Resolved by Invasion of a Novel Sex Determiner in Lake Malawi Cichlid Fishes. Science 326:998–1001.

Rousset, F. 2008. Genepop’007: A complete re-implementation of the Genepop software for Windows and Linux. Mol Ecol Resour 8:103–106.

Seeb, L. W., W. D. Templin, S. Sato, S. Abe, K. Warheit, J. Y. Park, and J. E. Seeb. 2011. Single nucleotide polymorphisms across a species’ range: implications for conservation studies of Pacific salmon. Mol Ecol Resour 11 Suppl 1:195–217.

Star, B., O. K. Tørresen, A. J. Nederbragt, K. S. Jakobsen, C. Pampoulie, and S. Jentoft. 2016. Genomic characterization of the Atlantic cod sex-locus. Scientific Reports 6:31235.

Tamate, T., and K. Maekawa. 2006. LATITUDINAL VARIATION IN SEXUAL SIZE DIMORPHISM OF SEA-RUN MASU SALMON, ONCORHYNCHUS MASOU. Evolution 60:196–201.

Thomas, J. W., M. Cáceres, J. J. Lowman, C. B. Morehouse, M. E. Short, E. L. Baldwin, D. L. Maney, and C. L. Martin. 2008. The Chromosomal Polymorphism Linked to Variation in Social Behavior in the White-Throated Sparrow (<em>Zonotrichia albicollis</em>) Is a Complex Rearrangement and Suppressor of Recombination. Genetics 179:1455–1468.

Tian, C., R. M. Plenge, M. Ransom, A. Lee, P. Villoslada, C. Selmi, L. Klareskog, A. E. Pulver, L. Qi, P. K. Gregersen, and M. F. Seldin. 2008. Analysis and Application of European Genetic Substructure Using 300 K SNP Information. PLoS Genet 4:e4.

Waples, R. K., L. W. Seeb, and J. E. Seeb. 2016. Linkage mapping with paralogs exposes regions of residual tetrasomic inheritance in chum salmon (Oncorhynchus keta). Mol Ecol Resour 16:17–28.

Wellenreuther, M., and L. Bernatchez. 2018. Eco-Evolutionary Genomics of Chromosomal Inversions. Trends Ecol Evol 33:427–440.

You, F. M., N. Huo, Y. Q. Gu, M. C. Luo, Y. Ma, D. Hane, G. R. Lazo, J. Dvorak, and O. D. Anderson. 2008. BatchPrimer3: a high throughput web application for PCR and sequencing primer design. BMC Bioinformatics 9:253.

